# A Systematic High-throughput Phenotyping Assay for Sugarcane Stalk Quality Characterization by Near-infrared Spectroscopy

**DOI:** 10.1101/2020.12.14.409383

**Authors:** Maoyao Wang, Xinru Li, Yinjuan Shen, Muhammad Adnan, Le Mao, Pan Lu, Qian Hu, Fuhong Jiang, Muhammad Tahir Khan, Zuhu Deng, Jiangfeng Huang, Muqing Zhang

## Abstract

Stalk quality improvement is deemed a promising strategy to enhance sugarcane production. However, the lack of efficient approaches for a systematic evaluation of sugarcane germplasm limited stalk quality improvement. In this study, 628 sugarcane samples were employed to take a high-throughput assay for determining the sugarcane stalk quality. Based on the high-performance anion chromatography method, large sugarcane stalk quality variations were detected in biomass composition and the corresponding fundamental ratio values. Online and offline Near-infrared Spectroscopy (NIRS) modeling strategies were applied for multiple purpose calibration. Consequently, 25 equations were generated with the excellent determination coefficient (*R^2^*) and ratio performance deviation (RPD) values. Notably, for some observations, RPD values as high as 6.3 were observed that indicated their exceptional performance potential and prediction capacity. Hence, this study provides a feasible way for high-throughput assessment of stalk quality, permitting large-scale screening of optimal sugarcane germplasm.

## 1. Introduction

Sugarcane (*Saccharum officinarum* L.) is a perennial C4 crop cultivated worldwide in the subtropical and tropical zones. It is one of the most important industrial crops for sugar and ethanol production (Moore et al., 2014). Moreover, sugarcane is an exceptionally productive commodity that is locally processed into value-added products and contributes to the cultivating areas’ economic welfare.

The sugarcane stalk quality plays a decisive role in the profitability of this crop. Sugar is the primary industrial product from the sugarcane stalk. Strenuous efforts have been made to obtain more sugar from cane stalks during the past decades. However, because of the complicated carbon partitioning and sugar accumulation mechanisms, limited achievements were gained (Bindon and Botha, 2002; Garcia Tavares et al., 2018; Ruan, 2014). Recently, advances in genomic tools and the decreasing costs of sequencing have enabled plant breeders to take large-scale precision breeding (Cavanagh et al., 2013; Watson et al., 2018). Despite such advances, the use of high-throughput phenotyping is anticipated to play a significant role in accelerating crop genetic improvement (Araus and Cairns, 2014; Furbank and Tester, 2011).

Sugarcane stalk consists of water, sugar, and fiber. These three significant components result in the dynamic variations of sugarcane stalk among different genotypes, growth periods, and meteorological conditions (Pereira et al., 2017). The dry mass of the cane stalks constituted by sugar and fiber, and therefore the sugar concentration is influenced by carbon partitioning between the two (Rohwer and Botha, 2001). The sugarcane payment is closely related to the sucrose concentration in fresh stalks determined by the stalk sucrose concentration on a dry weight basis (g sucrose g^-1^ DW) and moisture content (g water g^-1^ FW). Moreover, sugar composition and the ratio among different sugar forms also exhibit variation between different genotypes and within one genotype during sugarcane ripening (Chandra et al., 2015; Li et al., 2019). For instance, with the rapid accumulation of sucrose, the content of reducing sugars (glucose and fructose) gradually decreases as sugarcane matures. Hence, the ratio of reducing sugars to sucrose is usually used to evaluate the degree of sugarcane maturity (Verma et al., 2019). The precise analysis of these compounds in a high-throughput way may facilitate the large-scale accurate phenotyping of sugarcane stem quality.

Near-infrared spectroscopy (NIRS) has been applied as efficient tools for high-throughput screening of a large number of samples to predict their properties and compositions (Montes et al., 2007), especially for phenotyping and genomic selection in crop breeding (Cabrera-Bosquet et al., 2012; Ibraimo Samamad et al., 2018). It has been used for quality traits (such as juice soluble solids content, i.e., Brix, juice pH, firmness and water content) phenotyping in tomato (Ecarnot et al., 2013); estimation of sucrose, glucose, and fructose in sweet sorghum juice (Simeone et al., 2017); phenotyping of malt extract and protein content in barley (Walker et al., 2013); assessment of amino acid concentrations for QTL analysis in soybean (Warrington et al., 2015); quantitative monitoring of sucrose, reducing sugars and total sugar dynamics for phenotyping of water-deficit stress tolerance in rice (Das et al., 2018); prediction of silage quality traits for QTL mapping in maize (Seye et al., 2019); and herbage quality traits analysis (Cogan et al., 2005). Besides, NIRS has also been used for chemical compounds determination in sugarcane, such as for phosphorus analysis in leaves (Chen et al., 2002), mineral contents prediction under saline conditions (Steidle Neto et al., 2017), and cell wall components estimation in stalks (Hoang et al., 2017). Some studies have also attempted to employ NIRS calibration for sugar concentration in juice in terms of Brix or pol values (Hoang et al., 2017; Nawi et al., 2013) or commercial cane sugar contents (Sexton et al., 2018). However, little research has systematically explored NIRS assay for high-throughput characterization of sugarcane stalk quality.

In this work, hundreds of samples were collected from various genotypes at different growth stages. The stalk quality was assessed by quantitatively analyzing chemical composition and the corresponding ratio values in sugarcane stalk tissues upon high-performance anion chromatography (HPAEC-PAD) assay. Considerable variations of stalk quality were observed among these collections, allowing for a consistent offline and online NIRS calibration in sugarcane. This study provided systematic and multiple options-based assays for high-throughput screening of sugarcane biomass composition for stalk quality determination.

## 2. Materials and Methods

### 2.1. Sample collection

Sugarcane varieties were planted in the Fusui experimental field of Guangxi University, Nanning. The sugarcane stalks were collected once a month in five different periods from November 2018 to March 2019. After removing leaves and tips, six randomly selected stalks of each sugarcane variety were used for online NIRS spectrum scanning and further analysis. In addition, the stalks of forty sugarcane genotypies were collected every twenty days from the jointing stage to the ripening stage for model optimization.

### 2.2 Near-infrared spectral data collection

#### Online NIRS spectrum scanning

The randomly selected fresh stalks were immediately shredded using DM540 (IRBI Machines & Equipment Ltd, Brazil), blended and transmitted by CPS (Cane presentation system, Bruker Optik GmbH, Germany), and NIRS spectral data were simultaneously collected through the MATRIX-F (Bruker Optik GmbH, Germany) online system. The spectrum acquisition was taken by a full-band scanning mode at wavelengths ranging from 4000 to 10000 cm^-1^ with 4 cm^−1^ steps at room temperature. The spectral absorbance values were recorded as log1/R, where R is the sample reflectance. The obtained continuous reflectance values were then averaged for further analysis.

#### Offline NIRS spectrum scanning

The shredded fresh sugarcane samples were collected and dried under 60☐ after inactivation at 100 °C for one hour. The dried samples were ground over 40 mesh for offline NIRS spectrum data collection and further biomass chemical analysis. MATRIX-F equipped with a Q413 sensor head was used for contactless offline measurements. The reflectance of each sample was recorded and averaged for further calibration analysis.

### 2.3 Biomass component determination in sugarcane stalks

Moisture content was determined by a standard loss on drying method (Frazier and Westhoff, 1988). Sugar content (g/g, % dry weight) was analyzed by high-performance anion chromatography (HPAEC) method. Briefly, 0.100 g of ground dry sample was extracted with 40 mL ddH_2_O at 50☐ for 2 h. Additionally, 5.0 mL of lactose (1.0 mg/mL, Aladdin Biochemical Technology Co., Ltd., Shanghai, China) was added as an internal standard. The 50 mL sample was then filtered through 0.22 μm membrane filters for HPAEC detection.

ICS 5000^+^ system (Dionex/Thermo Fisher Scientific, Waltham, MA, USA) equipped with a pulsed amperometric detector (PAD) and Carbopac™ PA1 column (250 mm × 4 mm, 10 μm) was employed for determining soluble sugar in sugarcane. The chromatographic conditions were as following: column temperature was set at 30☐; injection volume was 25 μL; eluent A: ddH_2_O; and eluent B: 500 mmol/L NaOH solution (Merck KGaA, Darmstadt, Germany). An isocratic elution procedure of 60% A and 40% B at the flow rate of 2.0 mL/min was used for chromatographic analysis. The “Carbohydrates standard quad” waveform, as described in Annex 1, was employed for PAD.

For sugar content (g/g, % dry weight) calculation, the standard internal method was used for quantitative analysis. Analytical curves were produced using sucrose, D-glucose, and D-fructose as standards. Simultaneously, lactose was added as the internal standard (The standard chemicals were purchased from Aladdin Biochemical Technology Co., Ltd., Shanghai, China). The peak area ratios between each sugar (glucose, fructose, and sucrose) and the internal standard were calculated and corrected by the standard curves and then applied for its quantitative analysis. Insoluble residues content in sugarcane stalks was calculated by deducting the total soluble sugar from dry biomass. The biomass composition content (g/g, % fresh weight) was calculated based on its dry weight and the moisture content in fresh stalks. Biological triplicates were performed for each sample.

### 2.4 NIRS calibration

The OPUS spectroscopy software (version 7.8, Bruker Optik GmbH, Germany) was used for data processing and NIRS calibration. To solve the problems associated with overlapping peaks and baseline correction, pretreatments, and wavelength ranges, selection of the raw spectral data was performed before calibration. Several kinds of spectral pretreatment methods were provided in OPUS software, including constant offset elimination (COE), straight-line subtraction (SSL), standard normal variate (SNV), Min-Max normalization (MMN), multiplicative scattering correction (MSC), first derivative (FD), second derivative (SED), combinations of the first derivative and straight-line subtraction (FD+SSL), standard normal variate (FD+SNV), and multiplicative scattering correction (FD+MSC). The NIRS spectra were divided into multiple intervals and then reassembled to obtain the optimal spectral region. A principal component analysis (PCA) was conducted to characterize the structure of the spectral population, and the GH outlier (GH > 3.0) samples were eliminated. Partial least square (PLS) regression was performed to generate calibration equations. Internal cross-validation and external validation were carried out to test the performance of the generated calibration equations. The best equations were selected according to a high coefficient of determination of the calibration/internal cross-validation/external validation (*R^2^*/*R^2^cv*/*R^2^ev*), low root means square error of calibration/internal cross-validation/external validation (RMSEC/RMSECV/RMSEP), and high ratio performance deviation (RPD) values (Huang et al., 2012).

## 3. Results and Discussion

### 3.1 Precise sugar content determination in sugarcane stalks

The HPAEC-PAD assay was performed to detect sugar content in sugarcane stalks, and the standard internal method was adapted for quantitative analyses. In this assay system, all target compounds (glucose, fructose, and sucrose) and the internal standard (lactose) were separated entirely within 3.5 min (Fig. 1 A). Therefore, it allowed for rapid sugar content analysis in sugarcane stalks. The reducing sugars (glucose and fructose) content is supposed to be much lower than that of sucrose in the mature stem of sugarcane (Verma et al., 2019; Whittaker and Botha, 1997). To get more accurate equations for quantity analysis, a gap of 10 times difference in the concentrations was set between the sugars in gradient’s mixture for standard curve configuration. Expressly, the standard mixture for glucose and fructose was set to range from 0.25-8.0 μg/mL, while sucrose ranged from 2.5-80 μg/mL (Fig. 1B). As a result, high *R^2^* value for each sugar’s standard curves (glucose, fructose, and sucrose) was observed (Fig. 1C), which showed good performance of the quantitative analysis. In addition, to check whether the batch effects existed in this laboratory assay, the same sample was used for quantitative analysis of sugar content in different experimental batches. As shown in Fig. 1D, no significate differences were seen between them. Thus, the results indicated no batch effect in this quantitative analysis assay, suggesting that all the samples tested in the individual experimental batches could be combined for integrative analysis. Taken together, a rapid and stable HPAEC assay, allowing for an accurate analysis of sugar content in sugarcane stalks, was established.

**Fig. 1.**
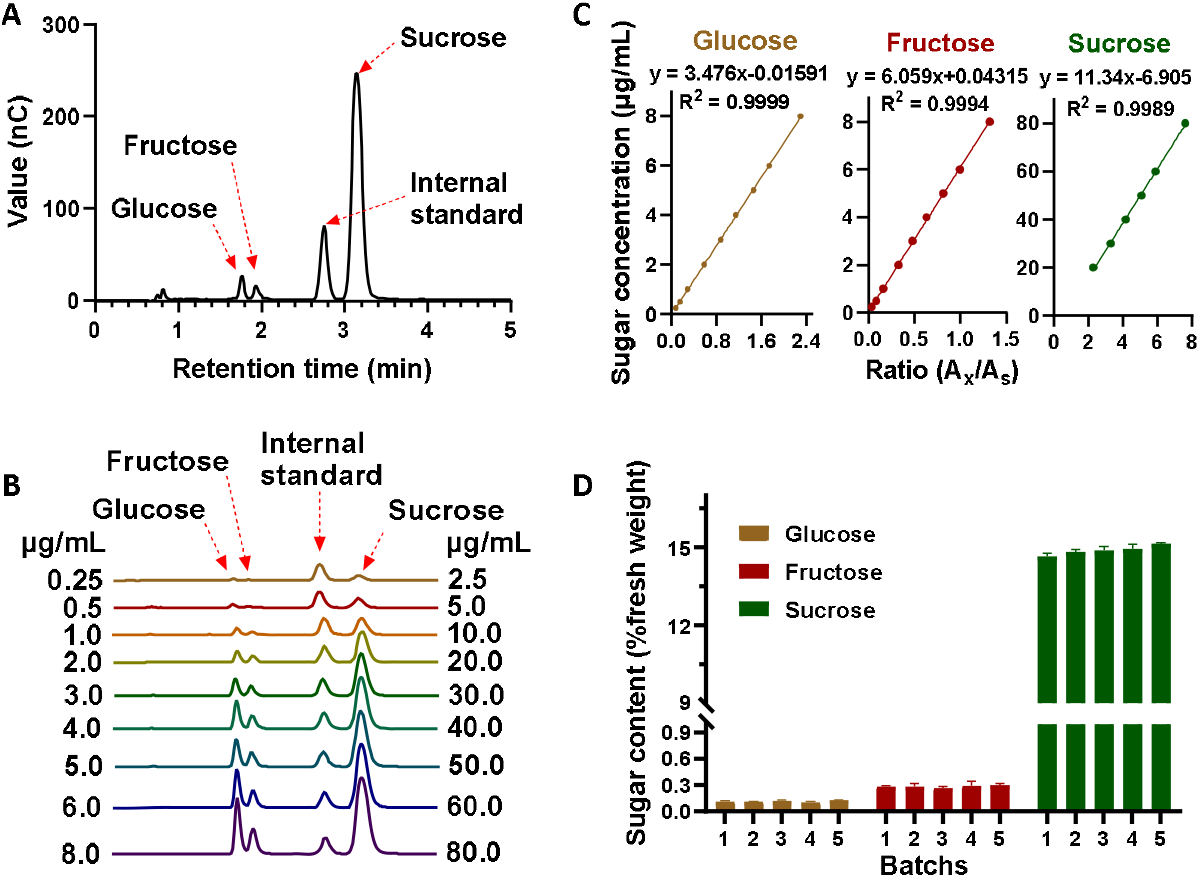
High performance anion chromatography assay for sugar determination in sugarcane. (A) chromatogram of sugar determination in sugarcane; (B) chromatogram of standard mixtures in different concentrations; (C) standard curves; (D) sugar determination in sugarcane at different batches.

### 3.2 Diverse biomass composition in collected sugarcane stalks

The biomass composition, especially for sugar content in cane stalks, is critical for its quality classification. To obtain samples with sufficient variability of biomass composition, sugarcane stalks of different genotypes were collected once a month from November 2018 to March 2019. Sugar mass content (g/g, % dry weight) of the ground dry samples were determined by the HPAEC-PAD assay described above. Due to genotype diversity, large variations were detected in each collection (Fig. 2A). Samples in collection 2 showed the highest diversity for reducing sugars, sucrose, and the total soluble sugars. Notably, a continuous increment of sucrose and total soluble sugar contents (g/g, % dry weight) was observed from collection 1 to 5 (Fig. 2A), which would have been due to the increasing maturity of the sugarcane stalks as time went on from November 2018 to March 2019. The different collections were combined to obtain a large sample set for NIRS calibration, as was shown in Fig. 2B. The integrated sample set exhibited a more comprehensive range of variation and better normal distribution. In detail, the reducing sugar content (g/g, % dry weight) ranged from 0.48 to 10.96 (average value at 2.87), sucrose ranged from 25.61 to 69.92, and the total soluble sugar content ranged from 27.02 to 73.88 (Fig. 2B). In sugarcane stem, soluble sugar and insoluble residues are central dry mass components formed due to photosynthesis. The insoluble residue was calculated by deducting total soluble sugar from dry biomass. Even though the insoluble residue content (g/g, % dry weight) in the collected sugarcane samples gradually decreased from collection 1 to 5 (Fig. 2C), a normal distribution ranging from 26.12 to 72.98 was observed in the combined sample set (Fig.2D).

**Fig. 2.**
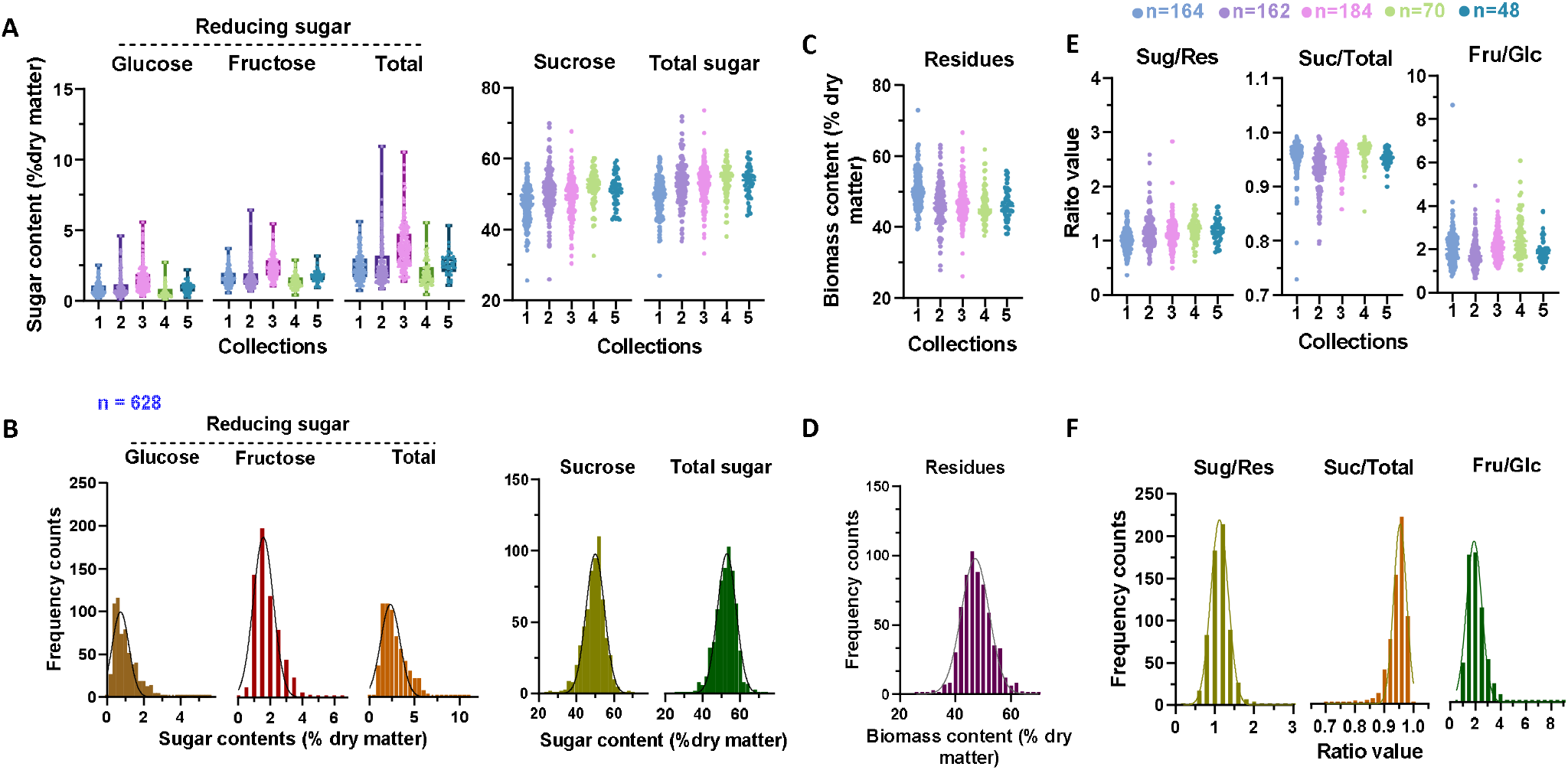
Variations of dry biomass composition in sugarcane stalks. Sugar mass content (A) and frequency distribution (B) in sugarcane stalks; insoluble residues content (C) and frequency distribution (D) in sugarcane stalks; ratio (E) between biomass composition and its frequency distribution (F). Varied genotypes of sugarcane collected at five different times, the number of each collections was 164, 162, 184, 70 and 48, respectively. Samples in different collections were merged together (n = 628) to calculate the distribution frequency of biomass component composition in B, D and F. Sug/Res, total soluble sugar/residues; Suc/Toal, sucrose/total soluble sugar; Fru/Glc, ratio of fructose/glucose in soluble sugar.

Generally, the proportion of chemical components in the sugarcane stalk is considered an important index to evaluate the quality (Lingle and Irvine, 1994). For example, the ratio between sugar and residues (Sug/Res) is closely related to the carbon partitioning patterns that primarily determine the clean sugar production in sugarcane stalks. In comparison, the sucrose proportion in total soluble sugar (Suc/Total) is recognized as the critical index for juice purity. The ratio between fructose/glucose (reducing sugars) relates to the physiological development of sugarcane (Lingle and Irvine, 1994; Thangavelu and Rao, 2002). These fundamental ratio values described above were calculated to carry out a systematic sugarcane stalk quality characterization. As expected, a considerable variation of these ratios was observed in the collected sugarcane population (Fig. 2E & F). Notably, the Sug/Res value ranged from 0.37 to 2.83, and a high coefficient of variation (CV) value was observed (0.23), suggesting a broad diversity of carbon partitioning patterns in the sugarcane population. In contrast, the Suc/Total value showed limited variation because most of the collected samples had almost matured during the study period.

For commercial sugarcane production, sugarcane stalk quality is determined by sucrose concentration on a fresh weight basis (g sucrose g^-1^ fresh weight). However, in fresh sugarcane stalk, the sugar concentration is not only related to the mass content of sugar in the dry matter but also close to the water content. The accumulation of % soluble sugars is reportedly associated with a concomitant decrease in moisture content (Lingle, 1999). An increase in sucrose content expressed in terms of the fresh mass may occur even without the deposition of additional sucrose when culms become dehydrated due to reduced soil moisture. Thus, sugarcane ripening reflected as sucrose % increases does not necessarily depict sucrose content (Glover, 1971). Therefore, in sugarcane, the high sugar mass content (g/g, % dry weight) and low moisture content (g/g, % fresh weight) could be considered an optimal sugar production source.

This study also determined the biomass concentration in fresh sugarcane stalk according to their dry biomass and moisture content. Owning to the classic drying water loss method (Frazier and Westhoff, 1988), moisture content diversity was detected in the collected sugarcane population (Fig. 3A & B). The water content of sugarcane decreased significantly from collection 1 to 5, which may be related to the gradual loss of water in the later stage of sugarcane growth (Fig. 3A). On the contrary, with the decrease in water content in sugarcane stalks, the sugar concentration (g/g, % fresh weight), mainly sucrose and total soluble sugar concentration, gradually increased (Fig. 3B). Whereas, the concentration of the residues (g/g, % fresh weight) seemed to be maintained at a stable level among the different collections (Fig. 3E). As expected, all of these compounds displayed a more considerable variation and showed a normal distribution in the combined sample set (Fig. 3B, D&F), which exhibited high NIRS calibration performance.

**Fig. 3.**
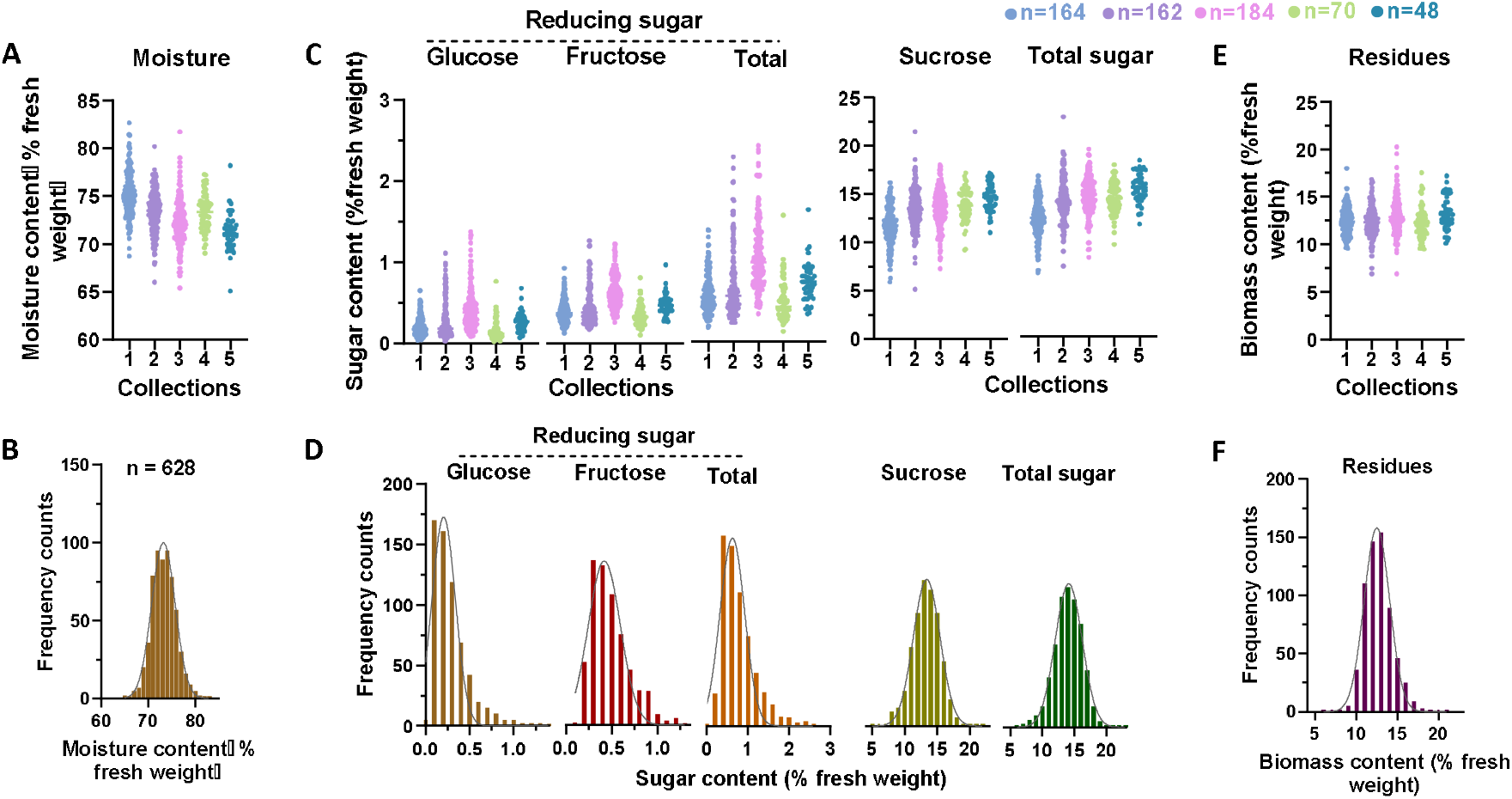
Variations of fresh biomass composition in sugarcane stalks. Moisture content (A) and frequency distribution (B) in sugarcane stalks; Sugar content (C) and frequency distribution (D) in sugarcane stalks; Insoluble residues content (E) and frequency distribution (F) in sugarcane stalks. Varied genotypes of sugarcane collected at five different times, the number of each collections was 164, 162, 184, 70 and 48, respectively. Samples in different collections were merged together (n=628) to calculate the distribution frequency of biomass component composition in B, D and F.

### 3.3 NIRS data characterization in collected sugarcane stalks

DM540-CPS coupling with the MATRIX-F system has been designed explicitly for sugarcane quality control (QC) analysis based on the related calibration model (Bruker, 2018). Sugarcane stalks were shredded and automatically passed to the NIR sensor for simultaneous spectral collection within one minute. Instead of pressing out the cane juice for quantity analysis, no juice was extracted from the sugarcane stalks. In order to establish the biomass composition calibration model for online quantity analysis, near-infrared spectroscopy of fresh sugarcane stalks for each collection was recorded on this system. As a result, broad diversity was detected in each sugarcane sample (Fig. 4A). PCA analysis was carried out to characterize the structure of the combined spectral population, as shown in Fig. 4B; no significate discrimination could be detected among these spectra from different collections. The continuous distribution of the combined spectral population further indicated that these samples could be integrated into a global NIRS calibration population. In addition, during the PCA analysis, global distance (GH) between each spectrum was calculated, while the GH outliers were eliminated from the population during further NIRS modeling.

**Fig. 4.**
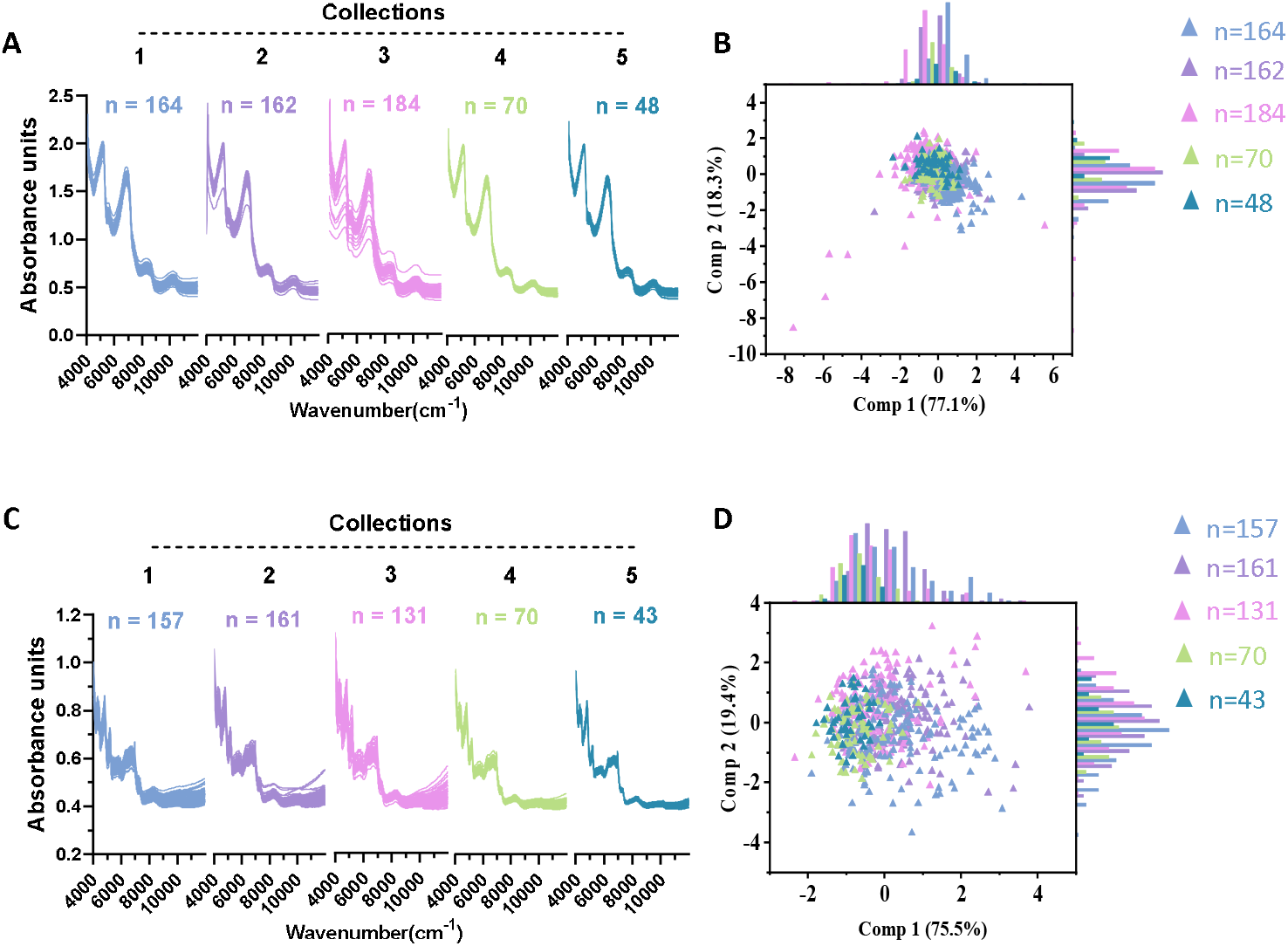
Variation of NIRS absorbance spectra in sugarcane samples. Original spectroscopy of fresh (A) and dry samples (C); PCA scores of near-infrared spectroscopy for fresh (B) and dry samples (D).

As a comparison, an offline near-infrared spectroscopy data scanning assay was applied to take the offline NIRS calibration. The dry ground samples from different collections were offline scanned by MATRIX-F equipped with a Q413 sensor. It was apparent from the results that the spectroscopy of the ground dry sample was different from that of the fresh sample (Fig. 4C), which can be attributed to the water loss. PCA results showed that the dry sample’s spectroscopy exhibited much higher variation (Fig. 4D), indicating that the offline assay’s spectra would be more conducive for NIRS modeling.

### 3.4 Determination of calibration and validation sets

A total of 562 samples in the combined sets were obtained for offline NIRS modeling. One in the fifth of the samples was randomly selected into the validation sets, while the remaining 449 samples formed the calibration sets. A descriptive statistical analysis was conducted to compare the calibration and validation sets in terms of the minimum (Min), maximum (Max), mean, standard deviation (SD), and coefficient of variation (CV) value (Table 1). Similarly, 628 samples were used for online NIRS modeling either for biomass composition content in dry weight (g/g, % dry weight) or in fresh sugarcane stalks (g/g, % fresh weight). Before NIRS modeling, 502 samples were randomly selected into the calibration sets, and the remaining 126 were included in the validation sets (Table 1). As shown in Table 1, all of the samples in the calibration and validation set showed comparable statistical distribution, allowing for reliable NIRS modeling.

**Table 1.** Calibration and validation sets for biomass components in sugarcane stalks.

|  | Calibration |  |  |  |  |  | Validation |  |  |  |  |  |
| --- | --- | --- | --- | --- | --- | --- | --- | --- | --- | --- | --- | --- |
|  | N<sup>a</sup> | Min<sup>b</sup> | Max<sup>c</sup> | Mean | SD<sup>d</sup> | CV<sup>e</sup> | N | Min | Max | Mean | SD | CV |
| **Dry weight (Offline)** |  |  |  |  |  |  |  |  |  |  |  |  |
| **Sugars** |  |  |  |  |  |  |  |  |  |  |  |  |
| Glucose | 449 | 0.10 | 4.76 | 0.98 | 0.69 | 0.71 | 113 | 0.22 | 5.55 | 1.29 | 0.99 | 0.77 |
| Fructose | 449 | 0.38 | 4.49 | 1.72 | 0.72 | 0.42 | 113 | 0.77 | 6.38 | 2.00 | 0.96 | 0.48 |
| Reducing sugar | 449 | 0.48 | 8.33 | 2.69 | 1.36 | 0.50 | 113 | 1.09 | 10.96 | 3.29 | 1.91 | 0.58 |
| Sucrose | 449 | 25.61 | 69.92 | 49.68 | 5.69 | 0.11 | 113 | 25.93 | 60.35 | 48.51 | 5.98 | 0.12 |
| Total sugar | 499 | 27.02 | 73.88 | 52.37 | 5.64 | 0.11 | 113 | 36.89 | 63.16 | 51.80 | 5.34 | 0.10 |
| **Residues** | 449 | 27.83 | 72.98 | 47.93 | 5.46 | 0.11 | 113 | 26.12 | 63.11 | 47.83 | 6.13 | 0.13 |
| **Ratio** |  |  |  |  |  |  |  |  |  |  |  |  |
| Sug/Res | 449 | 0.37 | 2.83 | 1.13 | 0.27 | 0.24 | 113 | 0.58 | 1.71 | 1.10 | 0.23 | 0.21 |
| Suc/Total | 449 | 0.81 | 0.99 | 0.95 | 0.03 | 0.03 | 113 | 0.70 | 0.98 | 0.94 | 0.04 | 0.05 |
| Fru/Glc | 449 | 0.67 | 8.65 | 2.12 | 0.78 | 0.37 | 113 | 0.74 | 4.03 | 1.89 | 0.65 | 0.34 |
| **Dry weight (Online)** |  |  |  |  |  |  |  |  |  |  |  |  |
| **Sugars** |  |  |  |  |  |  |  |  |  |  |  |  |
| Glucose | 502 | 0.10 | 5.12 | 1.04 | 0.72 | 0.69 | 126 | 0.23 | 5.55 | 1.14 | 0.94 | 0.82 |
| Fructose | 502 | 0.38 | 5.42 | 1.79 | 0.74 | 0.41 | 126 | 0.74 | 6.38 | 1.92 | 0.94 | 0.49 |
| Reducing sugar | 502 | 0.48 | 10.54 | 2.83 | 1.41 | 0.50 | 126 | 1.01 | 10.96 | 3.06 | 1.82 | 0.60 |
| Sucrose | 502 | 30.45 | 69.92 | 49.77 | 5.49 | 0.11 | 126 | 25.61 | 60.26 | 48.08 | 6.37 | 0.13 |
| Total sugar | 502 | 33.33 | 73.88 | 52.60 | 5.41 | 0.10 | 126 | 27.02 | 63.43 | 51.14 | 5.83 | 0.11 |
| **Residues** | 502 | 26.12 | 66.67 | 47.40 | 5.41 | 0.11 | 126 | 36.57 | 72.98 | 48.86 | 5.83 | 0.12 |
| **Ratio** |  |  |  |  |  |  |  |  |  |  |  |  |
| Sug/Res | 502 | 0.50 | 2.83 | 1.14 | 0.26 | 0.23 | 126 | 0.37 | 1.73 | 1.07 | 0.24 | 0.22 |
| Suc/Total | 502 | 0.79 | 0.99 | 0.95 | 0.03 | 0.03 | 126 | 0.70 | 0.98 | 0.94 | 0.04 | 0.04 |
| Fru/Glc | 502 | 0.69 | 8.65 | 2.07 | 0.78 | 0.34 | 126 | 0.67 | 3.97 | 2.04 | 0.63 | 0.31 |
| **Fresh weight (Online)** |  |  |  |  |  |  |  |  |  |  |  |  |
| **Moisture** | 502 | 65.10 | 81.74 | 73.30 | 2.43 | 0.03 | 126 | 65.40 | 82.68 | 73.52 | 3.12 | 0.04 |
| **Sugars** |  |  |  |  |  |  |  |  |  |  |  |  |
| Glucose | 502 | 0.03 | 1.34 | 0.28 | 0.20 | 0.71 | 126 | 0.06 | 1.38 | 0.30 | 0.24 | 0.81 |
| Fructose | 502 | 0.10 | 1.22 | 0.48 | 0.20 | 0.43 | 126 | 0.19 | 1.26 | 0.50 | 0.23 | 0.47 |
| Reducing sugar | 502 | 0.13 | 2.45 | 0.76 | 0.39 | 0.52 | 126 | 0.26 | 2.40 | 0.80 | 0.46 | 0.58 |
| Sucrose | 502 | 7.29 | 21.47 | 13.31 | 2.05 | 0.15 | 126 | 5.14 | 17.67 | 12.79 | 2.46 | 0.19 |
| Total sugar | 502 | 7.98 | 22.86 | 14.07 | 2.11 | 0.15 | 126 | 6.66 | 18.91 | 13.60 | 2.46 | 0.18 |
| **Residues** | 502 | 6.90 | 20.24 | 12.63 | 1.66 | 0.13 | 126 | 9.50 | 18.07 | 12.89 | 1.84 | 0.14 |

<sup>a</sup>N, sample number; <sup>b</sup>Min, minimum value; <sup>c</sup>Max, maximum value; <sup>d</sup>SD, standard deviation; <sup>e</sup>CV, coefficient of variation.

### 3.5 NIRS modeling for biomass compositions in sugarcane stalks

Partial least square (PLS) regression analysis methods packed in OPUS software were performed for NIRS modeling. The selected wavelength of near-infrared spectroscopy was pretreated with derivative and scatter correction methods before NIRS calibration. Internal cross-validation and external validation were applied to evaluate the calibration equations. During NIRS calibration, the root mean square error of calibration/cross-validation/ external validation (RMSEC/RMSECV/RMSEP), coefficient of determination of calibration/cross-validation/external validation (*R^2^*/*R^2^cv*/*R^2^ev*) and the ratio performance deviation (RPD) were obtained to select optima equations.

Due to the reduction of the water absorption in the near-infrared spectrum, offline NIRS calibration exhibits great advantage in dry biomass composition determination (Huang et al., 2017; Huang et al., 2012; Wu et al., 2015). In this study, the dry ground biomass of sugarcane stalks was applied for offline NIRS modeling. The results indicated that all of the equations for sugar, residues content (g/g, % dry weight), and the ratios obtained high *R^2^* value in calibration. This is especially true for sugar content (g/g, % dry weight) calibration, where the *R^2^* value reached as high as 0.91 (Annex 2). Besides, most of the equations exhibited high *R^2^cv* and RPD values during internal cross-validation, except for Fru/Glc, which showed a relatively low RPD value of 1.90 (Fig. 5 & Annex 2). Meanwhile, additional external validation was applied to evaluate the performance of these obtained equations. All of the equations exhibited a high linear correlation between the predicted and actual values. The glucose (g/g, % dry weight) showed the highest *R^2^cv* value of 0.92 (Fig. 5 & Annex 2). Notably, all of these equations showed the RPD value much higher than 2.0 during external validation (Fig. 5). The RPD values of equations were higher than 2.0 as promising ones (Li et al., 2017; Xu et al., 2013; Yang et al., 2016), and therefore, all of the equations obtained in this study for biomass composition (g/g, % dry weight) exhibited good prediction performance in offline NIRS detection system.

**Fig. 5.**
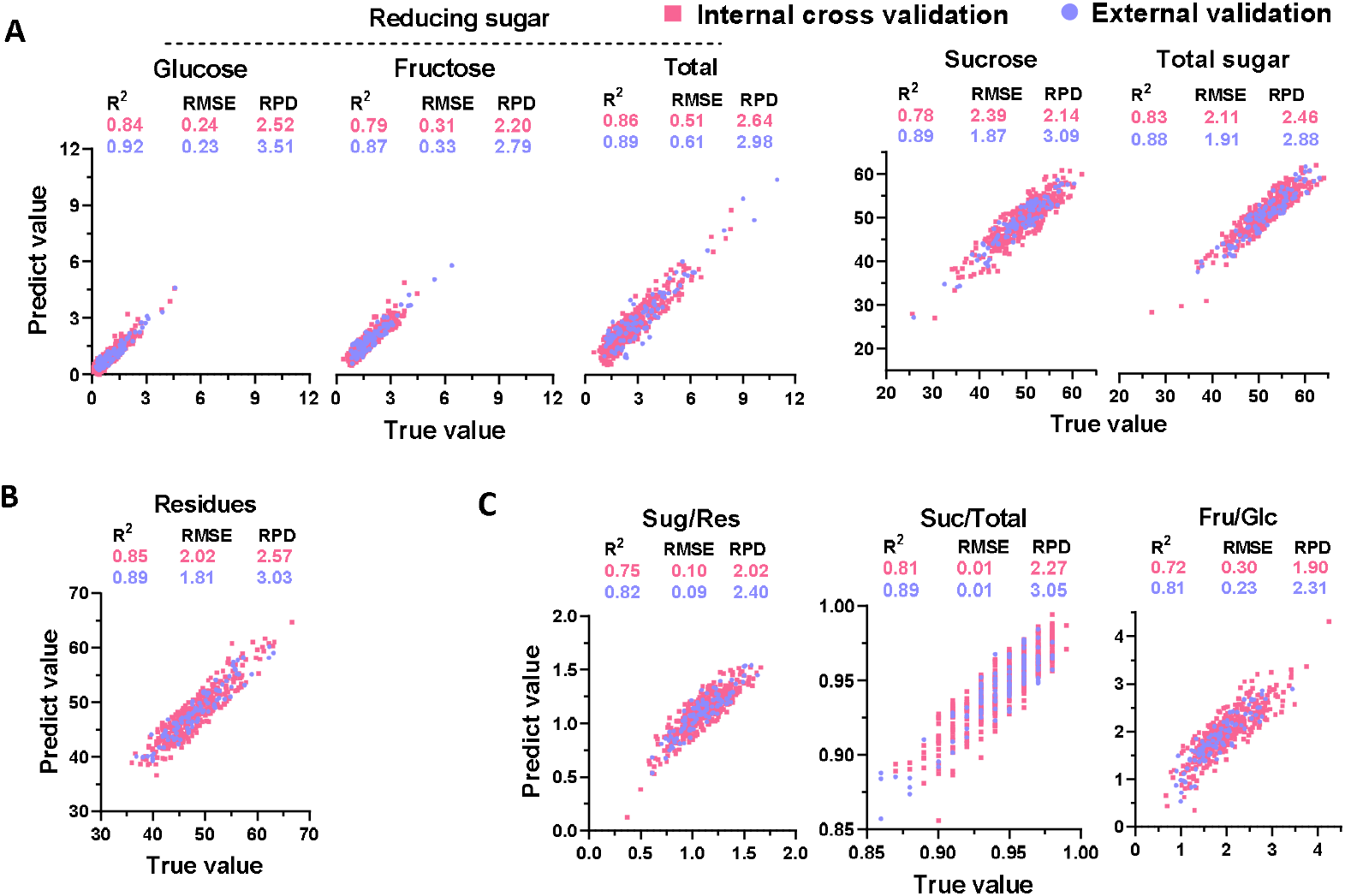
Correlation analysis between the predicted and true value for biomass component content (% dry matter) in sugarcane stalks upon offline NIRS calibration. A) sugar; B) insoluble residues; C) ratio between biomass component. The red and blue dots represented internal cross validation and external validation, respectively. *R^2^*, coefficient determination; RMSE, root mean square error; RPD, ratio performance deviation.

For comparison, online NIRS modeling was carried out for dry biomass composition content (g/g, % dry weight) prediction based on near-infrared spectra collected from fresh sugarcane stalks. The calibration results showed that, even though the equations obtained the *R^2^* values that were lower than the offline calibration, the results reached a substantially high level (ranging from 0.83 to 0.91) (Annex 2). Based on cross-validation and external validation data, most of the other equations obtained the RPD value over 2.0, except for reducing sugars (glucose, fructose, and the total, g/g, % dry weight), which showed relatively low *R^2^cv* ranging from 0.68 to 0.74 and RPD values ranging from 1.76 to 1.98 (Fig. 6). Notably, the ratio between sugar and residues (Sug/Res) exhibited the best performance in online NIRS calibration. It showed the highest *R^2^*, *R^2^cv*, and *R^2^ev* value of 0.91, 0.86, and 0.88, respectively (Fig. 6). As the ratio between sugar and residues (Sug/Res) was the key indicator of carbon partitioning pattern in sugarcane stalks, this study could provide a reliable high-throughput assay for large scale selection of promising germplasm from the sugarcane population through online NIRS system.

**Fig. 6.**
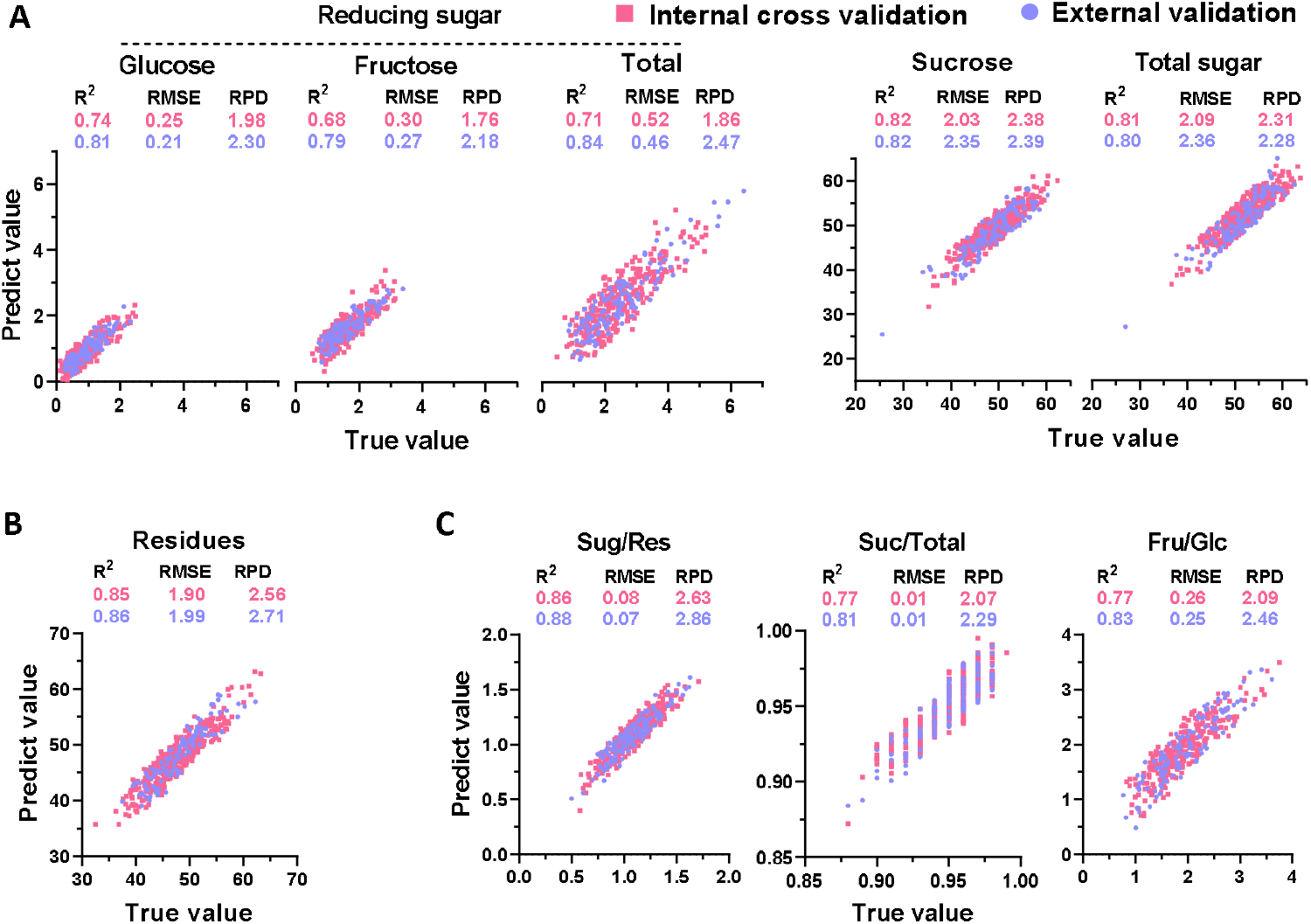
Correlation analysis between the predicted and true value for biomass component content (% dry matter) in sugarcane stalks upon online NIRS calibration. A) sugar; B) insoluble residues; C) ratio between biomass component. The red and blue dots represented internal cross validation and external validation, respectively. *R^2^*, coefficient determination; RMSE, root mean square error; RPD, ratio performance deviation.

Moreover, the near-infrared spectroscopy collected from fresh sugarcane stalks was applied for online NIRS calibration for biomass composition (g/g, % fresh weight) prediction. For sugar concentration (g/g, % fresh weight) modeling, the equations for sucrose and total soluble sugar exhibited the best performance with consistently higher *R^2^*, *R^2^cv*, *R^2^ev*, and RPD values during calibration and related validations (Fig. 7A & Annex 2). Meanwhile, the equations for reducing sugar concentration (g/g, % fresh weight) also exhibited consistently high RPD values over 2.0, indicating their excellent predictive capacity (Fig. 7A). In particular, the equation for moisture content (g/g, % fresh weight) showed a perfect linearly correlated relationship between the predicted and actual value, demonstrating a reliable and accurate online predictive capacity (Fig. 7B). Besides, the residues content (g/g, % fresh weight) also exhibited good predictive performance with consistent high *R^2^* value during calibration and two different kinds of validations (Fig. 7C).

**Fig. 7.**
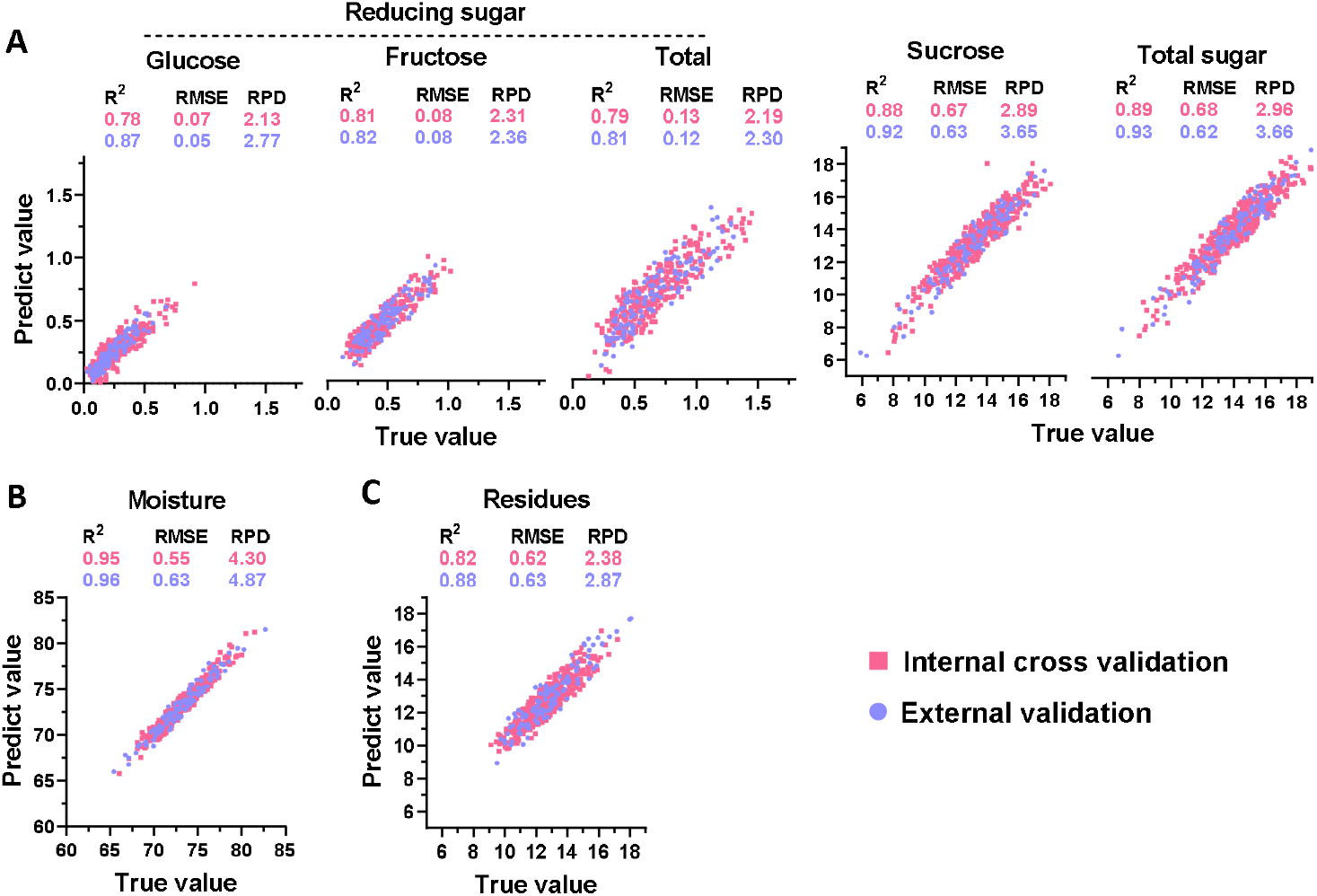
Correlation analysis between the predicted and true value for biomass component content (% fresh weight) in sugarcane stalks upon online NIRS calibration. A) sugar; B) moisture; C) insoluble residues. The red and blue dots represented internal cross validation and external validation, respectively. *R^2^*, coefficient determination; RMSE, root mean square error; RPD, ratio performance deviation.

Overall, it could be inferred from the data that, compared to the online and offline strategies for dry biomass composition modeling, the equations generated by offline calibration showed a higher prediction capacity (Fig. 5 & Fig. 6). When different sample types were compared during online NIRS modeling, the fresh sample’s biomass composition exhibited a much better performance (Fig. 6 & Fig. 7). Therefore, the data suggested that NIRS strategies could be considered to generate the optimal equations for a highly accurate prediction depending upon the sample type.

### 3.6 Integrative calibration for biomass composition in sugarcane stalks

To take a final NIRS calibration, samples in calibration and validation sets were integrated to form a final calibration set. Since more samples were contained in the final calibration set, all of the newly generated equations exhibited much better performance than those described above. In this approach, the average *R^2^* value increased from 0.88 to 0.93 by integrative calibration for offline prediction of dry biomass composition, and the average RPD value increased from 2.3 to 3.2 during cross-validation (Table 2 & Fig. 8A-C). For sugar content (g/g, % dry weight) offline prediction, all of the equations obtained high *R^2^* and *R^2^cv* values (over 0.90). The RPD values were higher than 3.0 for calibration and cross-validation (Table 2 & Fig. 8A). Thus, these equations exhibited excellent sugar content (g/g, % dry weight) determination via offline NIRS assay. The online NIRS modeling performance for dry biomass composition (g/g, % dry weight) did not improve as much as it bettered for offline NIRS modeling by expanding the calibration set. However, most of the equations obtained the RPD value over 2.0, permitting a reasonable prediction (Table 2 & Fig. 8 D-F). The integrative calibration processing enhanced the prediction capacity for online calibration of fresh biomass concentration (g/g, % fresh weight). Notably, apart from reducing sugars (g/g, % fresh weight) that showed the *R^2^* and *R^2^cv* values ranging from 0.82 to 0.93, all of the other equations obtained the *R^2^* and *R^2^cv* values much higher than 0.90, as well as high RPD values over 3.0 (Table 2 & Fig. 8G-I). Therefore, these newly generated equations could be applied for online quantitative analysis of biomass composition by NIRS assay.

**Table 2.** Integrative calibration statistics for optimized equations generated for prediction of biomass components in sugarcane stalks.

|  | Calibration |  |  |  |  |  |  |  | Cross validation |  |  |
| --- | --- | --- | --- | --- | --- | --- | --- | --- | --- | --- | --- |
|  | Rank | N | SCM | Spectrum range (cm<sup>-1</sup>) | Mean | SD | RMSEC | R<sup>2</sup> | RMSECV | R<sup>2</sup><sub>cv</sub> | RPD |
| **Dry weight (Offline)** |  |  |  |  |  |  |  |  |  |  |  |
| **Sugars (% dry weigh)** |  |  |  |  |  |  |  |  |  |  |  |
| Glucose | 15 | 464 | FD | 3996~11988 | 0.96 | 0.66 | 0.15 | 0.95 | 0.19 | 0.92 | 3.44 |
| Fructose | 16 | 418 | FD | 3996~11988 | 1.70 | 0.75 | 0.16 | 0.96 | 0.22 | 0.92 | 3.44 |
| Reducing sugar | 15 | 457 | FD | 3996~11988 | 2.74 | 1.48 | 0.33 | 0.95 | 0.42 | 0.92 | 3.50 |
| Sucrose | 10 | 394 | FD+SNV | 3996~11988 | 48.93 | 5.17 | 1.29 | 0.94 | 1.49 | 0.92 | 3.48 |
| Total sugar | 10 | 410 | FD+SNV | 3996~11988 | 51.85 | 5.27 | 1.32 | 0.94 | 1.51 | 0.92 | 3.49 |
| **Residues** | 8 | 410 | FD | 4759.7~9411.4 | 48.16 | 5.34 | 1.59 | 0.91 | 1.68 | 0.90 | 3.18 |
| **Ratio** |  |  |  |  |  |  |  |  |  |  |  |
| Sug/Res | 10 | 425 | FD | 3996~7197.4, 1185.7~11988 | 1.08 | 0.21 | 0.07 | 0.90 | 0.07 | 0.88 | 2.94 |
| Suc/Total | 13 | 536 | SNV | 4242.8~8894.5 | 0.95 | 0.03 | 0.01 | 0.87 | 0.01 | 0.83 | 2.45 |
| Fru/Glc | 13 | 427 | FD | 4104~7251.4, 8825.1~11185.7 | 1.95 | 0.57 | 0.17 | 0.92 | 0.21 | 0.87 | 2.75 |
| **Dry weight (Online)** |  |  |  |  |  |  |  |  |  |  |  |
| **Sugars** |  |  |  |  |  |  |  |  |  |  |  |
| Glucose | 19 | 407 | FD | 4890.8~11185.7 | 0.89 | 0.49 | 0.18 | 0.88 | 0.21 | 0.82 | 2.33 |
| Fructose | 12 | 391 | FD | 4890.8~7251.4, 8038.3~11185.7 | 1.63 | 0.57 | 0.27 | 0.78 | 0.29 | 0.74 | 1.98 |
| Reducing sugar | 22 | 401 | FD | 4242.8~4767.4, 5785.7~9411.4 | 2.46 | 1.02 | 0.38 | 0.87 | 0.44 | 0.81 | 2.31 |
| Sucrose | 15 | 514 | FD | 4890.8~11972.5 | 49.52 | 4.92 | 1.80 | 0.87 | 1.98 | 0.83 | 2.49 |
| Total sugar | 13 | 514 | FD | 4890.8~11185.7 | 52.30 | 5.01 | 1.84 | 0.87 | 1.96 | 0.85 | 2.56 |
| **Residues** | 12 | 495 | FD | 4890.8~8825.1, 9612~11972.5 | 47.52 | 5.01 | 1.75 | 0.88 | 1.86 | 0.86 | 2.69 |
| **Ratio** |  |  |  |  |  |  |  |  |  |  |  |
| Sug/Res | 24 | 473 | FD | 4104~11972.5 | 1.10 | 0.20 | 0.06 | 0.91 | 0.08 | 0.86 | 2.70 |
| Suc/Total | 18 | 396 | FD+SNV | 4752~6834.8, 7344~9411.4 | 0.95 | 0.02 | 0.01 | 0.86 | 0.01 | 0.81 | 2.27 |
| Fru/Glc | 31 | 396 | FD | 4104~11972.5 | 1.94 | 0.56 | 0.18 | 0.91 | 0.23 | 0.83 | 2.41 |
| **Fresh weight (Online)** |  |  |  |  |  |  |  |  |  |  |  |
| **Moisture** | 23 | 528 | FD+SNV | 4104~11972.5 | 73.21 | 2.46 | 0.40 | 0.97 | 0.46 | 0.97 | 5.40 |
| **Sugars** |  |  |  |  |  |  |  |  |  |  |  |
| Glucose | 34 | 415 | FD | 4104~11972.5 | 0.25 | 0.15 | 0.04 | 0.93 | 0.06 | 0.86 | 2.66 |
| Fructose | 18 | 405 | FD+SNV | 4104~11972.5 | 0.46 | 0.18 | 0.07 | 0.86 | 0.08 | 0.82 | 2.36 |
| Reducing sugar | 24 | 403 | FD | 4242.8~5276.6, 5785.7~9411.4 | 0.67 | 0.28 | 0.10 | 0.88 | 0.12 | 0.83 | 2.39 |
| Sucrose | 13 | 453 | FD+SSL | 4782.8~11185.7 | 13.27 | 1.91 | 0.45 | 0.95 | 0.49 | 0.93 | 3.88 |
| Total sugar | 23 | 514 | FD | 4104~11972.5 | 13.97 | 2.09 | 0.50 | 0.95 | 0.57 | 0.93 | 3.68 |
| **Residues** | 23 | 459 | FD | 4104~11972.5 | 12.66 | 1.55 | 0.43 | 0.93 | 0.48 | 0.90 | 3.21 |

N, sample number; SCM, scatter correction methods; SD, standard deviation of reference value; RMSEC, root mean square error of calibration; R<sup>2</sup>, determination coefficient; RMSECV, root mean square err of cross validation; R<sup>2</sup><sub>cv</sub>, determination coefficient of cross validation; RPD, ratio performance deviation; SNV, standard normal variate; SSL, straight line subtraction; FD, first derivative; FD+SNV, a combination of FD and SNV; FD+SSL, a combination of FD and SSL.

**Fig. 8.**
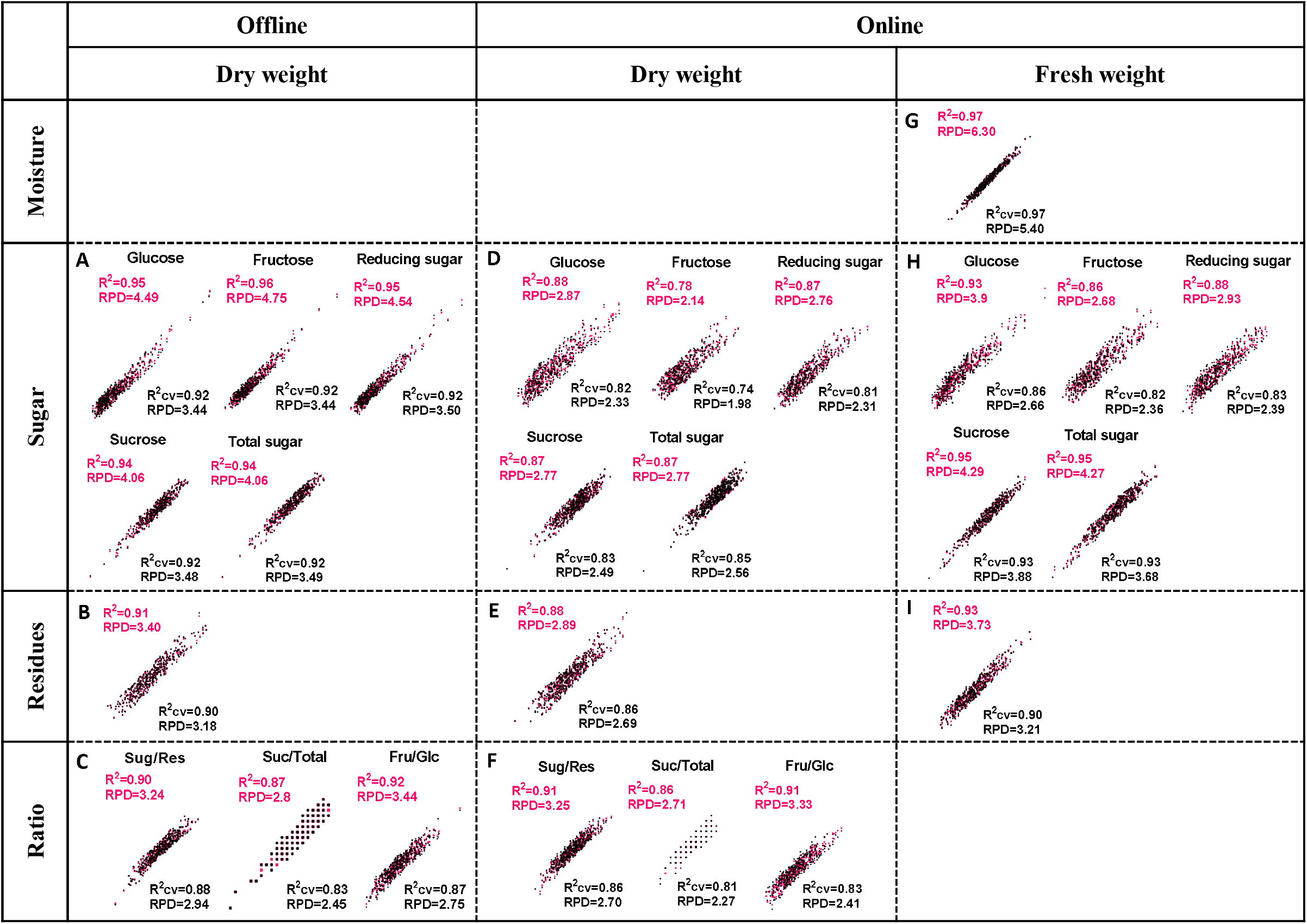
Correlation analysis between the fit (predict) and true value for biomass component content in sugarcane stalks. A-C, offline NIRS calibration for dry biomass of sugarcane stalks upon sugar content (A), residues (B) and ratio between them (C); D-F online NIRS calibration for dry biomass of sugarcane stalks on sugar content (D), residues (E) and the ratio (F); G-I, on-line NIRS calibration for fresh biomass of sugarcane stalks upon moisture content (G), sugar content (H) and residues (I). The red and black color represented calibration and internal cross validation, respectively.

A considerable improvement in prediction capacity was observed in both offline and online NIRS modeling via integrative calibration. The newly generated equations should be applicable for the biomass composition content prediction. The suggested models provide multiple options for the related high-throughput screen jobs. Notably, the online calibration models can play a significant role as they are substantially advantageous in high-throughput analysis of large-scale samples and offer better prospects for practical applications in the future.

## 4. Conclusions

A total of 628 sugarcane accessions were collected from different maturing stages of sugarcane stalk development. HPACE-PAD detected considerable variations of dry biomass composition and the related parameters in these collections’ fresh stalks. Offline and online NIRS modeling was carried out to take a systematic NIRS calibration. Twenty-five equations were generated with high *R^2^*, *R^2^cv*, *R^2^ev*, and RPD values, allowing for a consistent NIRS assay for multipurpose biomass composition predictions in sugarcane. Hence, this study provided a high-throughput strategy for large-scale screening of optimal sugarcane varieties and precision breeding.

## Supporting information

Electronic A. Supplementary data

## Acknowledgments

This work was funded by the Science and Technology Major Project of Guangxi (AA17202042-7 and Gui Ke 2018-266-Z01); the State Key Laboratory for Conservation and Utilization of Subtropical Agro-Bioresources (SKLCUSA-a202002).

## Electronic A. Supplementary data

Supplementary data associated with this article can be found in the online version.

**Annex 1**Waveform used in PAD for HPAEC detecting

**Annex 2**Statistics for equations generated for prediction of biomass components in sugarcane stalks.

## References

Araus, J.L., Cairns, J.E., 2014. Field high-throughput phenotyping: the new crop breeding frontier. Trends Plant Sci 19, 52–61.

Bindon, K.A., Botha, F.C., 2002. Carbon allocation to the insoluble fraction, respiration and triose-phosphate cycling in the sugarcane culm. Physiologia Plantarum 116, 12–19.

Bruker, 2018. FT-NIR analyzers for QC in the lab and production. Available at: https://www.bruker.com/fileadmin/user_upload/8-PDF-Docs/OpticalSpectrospcopy/FT-NIR/MATRIX-F/Brochures/Sugar_NIR_Brochure_EN.pdf Accessed 7 Sep 2020.

Cabrera-Bosquet, L., Crossa, J., von Zitzewitz, J., Dolors Serret, Serret., Luis Araus, Araus., 2012. High-throughput phenotyping and genomic selection: The frontiers of crop breeding converge. J Integr Plant Biol 54, 312–320.

Cavanagh, C.R., Chao, S., Wang, S., Huang, B.E., Stephen, S., Kiani, S., Forrest, K., Saintenac, C., Brown-Guedira, G.L., Akhunova, A., See, D., Bai, G., Pumphrey, M., Tomar, L., Wong, D., Kong, S., Reynolds, M., da Silva, M.L., Bockelman, H., Talbert, L., Anderson, J.A., Dreisigacker, S., Baenziger, S., Carter, A., Korzun, V., Morrell, P.L., Dubcovsky, J., Morell, M.K., Sorrells, M.E., Hayden, M.J., Akhunov, E., 2013. Genome-wide comparative diversity uncovers multiple targets of selection for improvement in hexaploid wheat landraces and cultivars. Proc Natl Acad Sci U S A 110, 8057–8062.

Chandra, A., Verma, P.K., Islam, M.N., Grisham, M.P., Jain, R., Sharma, A., Roopendra, K., Singh, K., Singh, P., Verma, I., Solomon, S., 2015. Expression analysis of genes associated with sucrose accumulation in sugarcane (Saccharum spp. hybrids) varieties differing in content and time of peak sucrose storage. Plant Biol (Stuttg) 17, 608–617.

Chen, M., Glaz, B., Gilbert, R.A., Daroub, S.H., Barton, F.E., Wan, Y., 2002. Near-infrared reflectance spectroscopy analysis of phosphorus in sugarcane leaves. Agron J 94, 1324–1331.

Cogan, N., Smith, K., Yamada, T., Francki, M., Vecchies, A., Jones, E., Spangenberg, G., Forster, J., 2005. QTL analysis and comparative genomics of herbage quality traits in perennial ryegrass (Lolium perenne L.). Theor Appl Genet 110, 364–380.

Das, B., Sahoo, R.N., Pargal, S., Krishna, G., Verma, R., Chinnusamy, V., Sehgal, V.K., Gupta, V.K., Dash, S.K., Swain, P., 2018. Quantitative monitoring of sucrose, reducing sugar and total sugar dynamics for phenotyping of water-deficit stress tolerance in rice through spectroscopy and chemometrics. Spectroc Acta Pt A-Molec Biomolec Spectr 192, 41–51.

Ecarnot, M., Baczyk, P., Tessarotto, L., Chervin, C., 2013. Rapid phenotyping of the tomato fruit model, Micro-Tom, with a portable VIS-NIR spectrometer. Plant Physiol Bioch 70, 159–163.

Frazier, W.C., Westhoff, D.C., 1988. Official Methods of Analysis of AOAC International. 6th ed. Volume 11 AOAC International Publishers; Gaithersburg, MA, USA: 1999. Food Microbiology 4th ed; International Edition; McGraw-Hill: Singapore, 1988; 440p.AOAC. 5th Revision.

Furbank, R.T., Tester, M., 2011. Phenomics - technologies to relieve the phenotyping bottleneck. Trends Plant Sci 16, 635–644.

Garcia Tavares, R., Lakshmanan, P., Peiter, E., O’Connell, A., Caldana, C., Vicentini, R., Soares, J.S., Menossi, M., 2018. ScGAI is a key regulator of culm development in sugarcane. J Exp Bot 69, 3823–3837.

Glover, J., 1971. Changes in sucrose % cane and yield of sucrose per unit area associated with cold, drought and ripening. Proceedings of the South African Sugar Technologists’ Association, 158–164.

Hoang, N.V., Furtado, A., Donnan, L., Keeffe, E.C., Botha, F.C., Henry, R.J., 2017. High-throughput profiling of the fiber and sugar composition of sugarcane biomass. Bioenerg Res 10, 400–416.

Huang, J., Li, Y., Wang, Y., Chen, Y., Liu, M., Wang, Y., Zhang, R., Zhou, S., Li, J., Tu, Y., Hao, B., Peng, L., Xia, T., 2017. A precise and consistent assay for major wall polymer features that distinctively determine biomass saccharification in transgenic rice by near-infrared spectroscopy. Biotechnol Biofuels 10.

Huang, J., Xia, T., Li, A., Yu, B., Li, Q., Tu, Y., Zhang, W., Yi, Z., Peng, L., 2012. A rapid and consistent near infrared spectroscopic assay for biomass enzymatic digestibility upon various physical and chemical pretreatments in Miscanthus. Bioresour Technol 121, 274–281.

Ibraimo Samamad, N.T., Dias Ribeiro, L.P., de Almeida Lopes, M.M., Puschmann, R., Silva, E.d.O., 2018. Near infrared spectroscopy, a suitable tool for fast phenotyping - The case of cashew genetic improvement. Sci Horti 238, 363–368.

Li, M., Wang, J., Du, F., Diallo, B., Xie, G.H., 2017. High-throughput analysis of chemical components and theoretical ethanol yield of dedicated bioenergy sorghum using dual-optimized partial least squares calibration models. Biotechnol Biofuels 10, 206.

Li, Y., Wang, W., Feng, Y., Tu, M., Wittich, P.E., Bate, N.J., Messing, J., 2019. Transcriptome and metabolome reveal distinct carbon allocation patterns during internode sugar accumulation in different sorghum genotypes. Plant Biotechnol J 17, 472–487.

Lingle, S.E., 1999. Sugar metabolism during growth and development in sugarcane internodes. Crop Sci 39, 480–486.

Lingle, S.E., Irvine, J.E., 1994. Sucrose synthase and natural ripening in sugarcane. Crop Sci 34, 1279–1283.

Montes, J.M., Melchinger, A.E., Reif, J.C., 2007. Novel throughput phenotyping platforms in plant genetic studies. Trends Plant Sci 12, 433–436.

Moore, P.H., Paterson, A.H., Tew, T., 2014. Sugarcane: The crop, the plant, and domestication, in: Moore, P.H., Botha, F.C. (Eds.), Sugarcane: physiology, biochemistry, and functional biology. Wiley-Blackwell Publishing, New Jersey, pp. 623–643.

Nawi, N.M., Chen, G., Jensen, T., Mehdizadeh, S.A., 2013. Prediction and classification of sugar content of sugarcane based on skin scanning using visible and shortwave near infrared. Biosyst Eng 115, 154–161.

Pereira, L.F.M., Ferreira, V.M., Oliveira, N.G., Sarmento, P., Endres, L., Teodoro, I., 2017. Sugars levels of four sugarcane genotypes in different stem portions during the maturation phase. An Acad Bras Cienc 89, 1231–1242.

Rohwer, J.M., Botha, F.C., 2001. Analysis of sucrose accumulation in the sugar cane culm on the basis of in vitro kinetic data. Biochemical Society 358, 437–445.

Ruan, Y.L., 2014. Sucrose metabolism: gateway to diverse carbon use and sugar signaling. Annu Rev Plant Biol 65, 33–67.

Sexton, J., Everingham, Y., Donald, D., Staunton, S., White, R., 2018. A comparison of non-linear regression methods for improved on-line near infrared spectroscopic analysis of a sugarcane quality measure. J Near Infrared Spec 26, 297–310.

Seye, A.I., Bauland, C., Giraud, H., Mechin, V., Reymond, M., Charcosset, A., Moreau, L., 2019. Quantitative trait loci mapping in hybrids between Dent and Flint maize multiparental populations reveals group-specific QTL for silage quality traits with variable pleiotropic effects on yield. Theor Appl Genet 132, 1523–1542.

Simeone, M.L.F., Parrella, R.A.C., Schaffert, R.E., Damasceno, C.M.B., Leal, M.C.B., Pasquini, C., 2017. Near infrared spectroscopy determination of sucrose, glucose and fructose in sweet sorghum juice. Microchem J 134, 125–130.

Steidle Neto, A.J., Toledo, J.V., Zolnier, S., Lopes, D.d.C., Pires, C.V., da Silva, T.G.F., 2017. Prediction of mineral contents in sugarcane cultivated under saline conditions based on stalk scanning by Vis/NIR spectral reflectance. Biosyst Eng 156, 17–26.

Thangavelu, S., Rao, K.C., 2002. Fructose-glucose ratio - A method to identify and classify the maturity of sugarcane. Sugar Tech 4, 66–68.

Verma, I., Roopendra, K., Sharma, A., Chandra, A., Kamal, A., 2019. Expression analysis of genes associated with sucrose accumulation and its effect on source-sink relationship in high sucrose accumulating early maturing sugarcane variety. Physiol Mol Biol Plants 25, 207–220.

Walker, C.K., Ford, R., Munoz-Amatriain, M., Panozzo, J.F., 2013. The detection of QTLs in barley associated with endosperm hardness, grain density, grain size and malting quality using rapid phenotyping tools. Theor Appl Genet 126, 2533–2551.

Warrington, C.V., Abdel-Haleem, H., Hyten, D.L., Cregan, P.B., Orf, J.H., Killam, A.S., Bajjalieh, N., Li, Z., Boerma, H.R., 2015. QTL for seed protein and amino acids in the Benning x Danbaekkong soybean population. Theor Appl Genet 128, 839–850.

Watson, A., Ghosh, S., Williams, M.J., Cuddy, W.S., Simmonds, J., Rey, M.D., Hatta, M.A.M., Hinchliffe, A., Steed, A., Reynolds, D., Adamski, N.M., Breakspear, A., Korolev, A., Rayner, T., Dixon, L.E., Riaz, A., Martin, W., Ryan, M., Edwards, D., Batley, J., Raman, H., Carter, J., Rogers, C., Domoney, C., Moore, G., Harwood, W., Nicholson, P., Dieters, M.J., DeLacy, I.H., Zhou, J., Uauy, C., Boden, S.A., Park, R.F., Wulff, B.B.H., Hickey, L.T., 2018. Speed breeding is a powerful tool to accelerate crop research and breeding. Nat Plants 4, 23–29.

Whittaker, A., Botha, F.C., 1997. Carbon partitioning during sucrose accumulation in sugarcane internodal tissue. Plant Physiol 115, 1651–1651 1659.

Wu, L.M., Li, M., Huang, J.F., Zhang, H., Zou, W.H., Hu, S.W., Li, Y., Fan, C.F., Zhang, R., Jing, H.C., Peng, L.C., Feng, S.Q., 2015. A near infrared spectroscopic assay for stalk soluble sugars, bagasse enzymatic saccharification and wall polymers in sweet sorghum. Bioresour Technol 177, 118–124.

Xu, F., Yu, J., Tesso, T., Dowell, F., Wang, D., 2013. Qualitative and quantitative analysis of lignocellulosic biomass using infrared techniques: A mini-review. Applied Energy 104, 801–809.

Yang, Z., Li, K., Zhang, M., Xin, D., Zhang, J., 2016. Rapid determination of chemical composition and classification of bamboo fractions using visible-near infrared spectroscopy coupled with multivariate data analysis. Biotechnol Biofuels 9, 35.

