## Supplementary material for "A Systematic High-throughput Phenotyping Assay for Sugarcane Stalk Quality Characterization by Near-infrared Spectroscopy": Electronic A. Supplementary data

### Slide 1
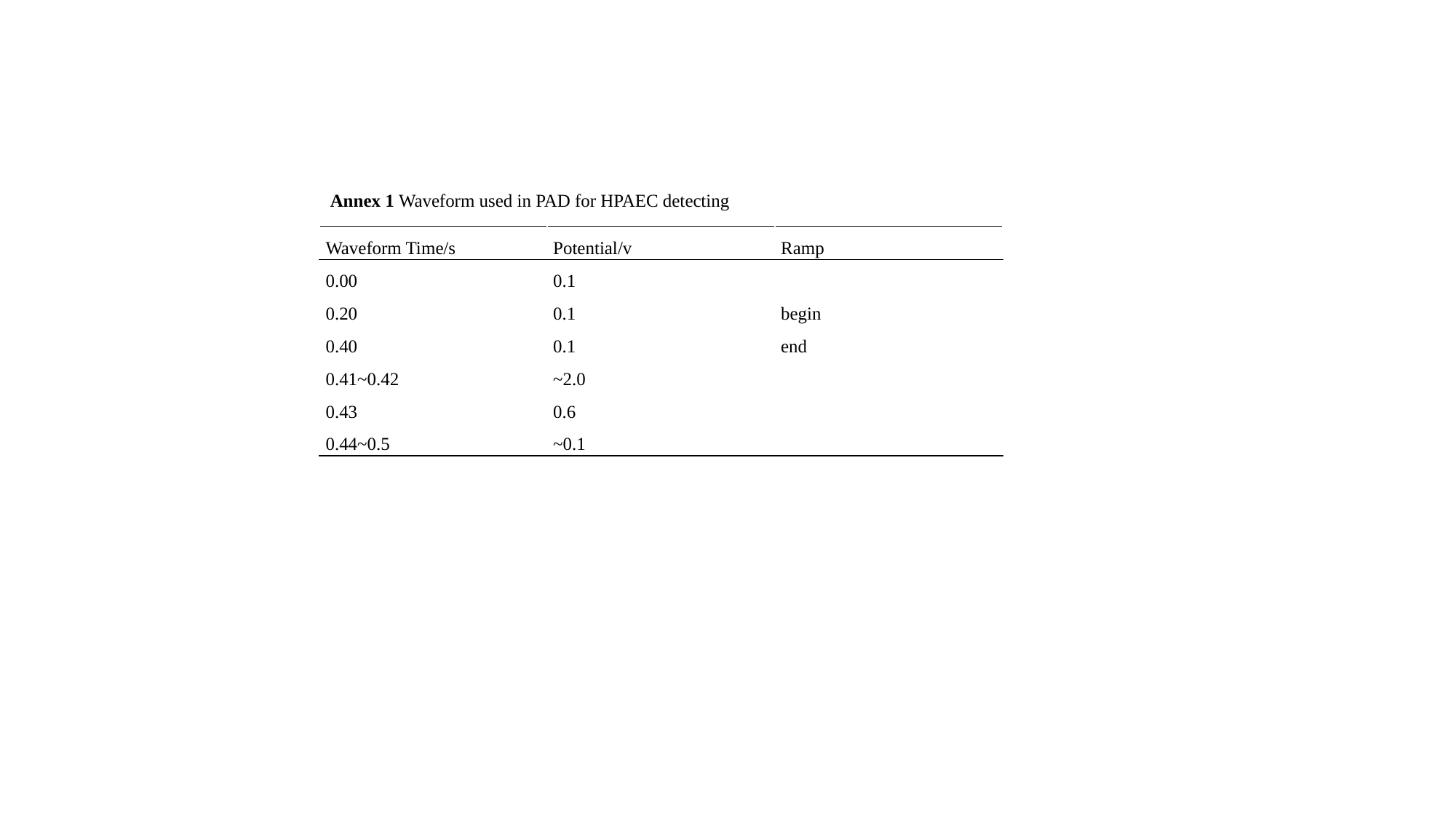

Annex 1 Waveform used in PAD for HPAEC detecting
| Waveform Time/s | Potential/v | Ramp |
| --- | --- | --- |
| 0.00 | 0.1 | |
| 0.20 | 0.1 | begin |
| 0.40 | 0.1 | end |
| 0.41~0.42 | ~2.0 | |
| 0.43 | 0.6 | |
| 0.44~0.5 | ~0.1 | |

### Slide 2
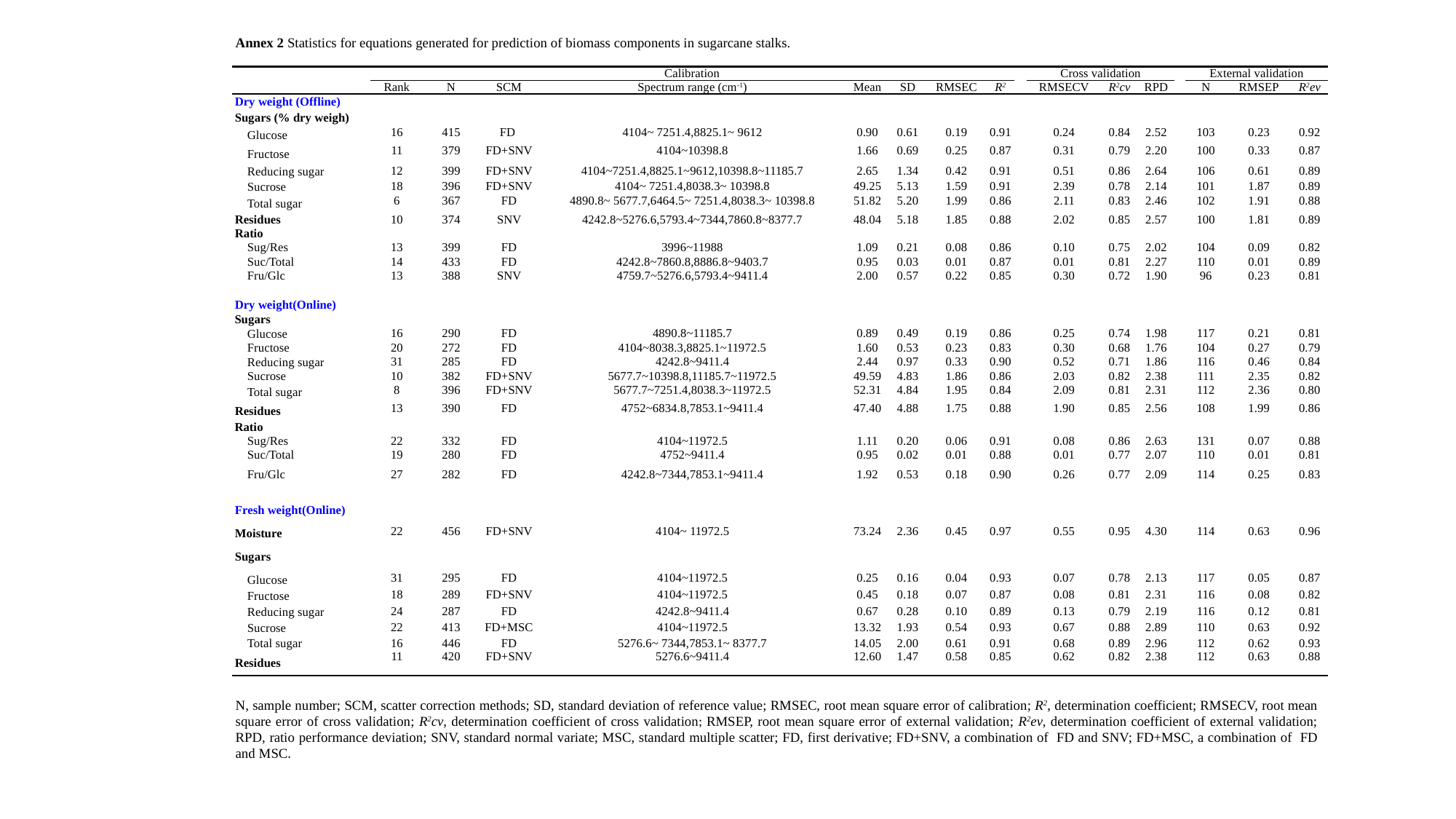

Annex 2 Statistics for equations generated for prediction of biomass components in sugarcane stalks.
| | Calibration | | | | | | | | | Cross validation | | | | External validation | | |
| --- | --- | --- | --- | --- | --- | --- | --- | --- | --- | --- | --- | --- | --- | --- | --- | --- |
| | Rank | N | SCM | Spectrum range (cm^-1^) | Mean | SD | RMSEC | R^2^ | | RMSECV | R^2^cv | RPD | | N | RMSEP | R^2^ev |
| Dry weight (Offline) | | | | | | | | | | | | | | | | |
| Sugars (% dry weigh) | | | | | | | | | | | | | | | | |
| Glucose | 16 | 415 | FD | 4104~ 7251.4,8825.1~ 9612 | 0.90 | 0.61 | 0.19 | 0.91 | | 0.24 | 0.84 | 2.52 | | 103 | 0.23 | 0.92 |
| Fructose | 11 | 379 | FD+SNV | 4104~10398.8 | 1.66 | 0.69 | 0.25 | 0.87 | | 0.31 | 0.79 | 2.20 | | 100 | 0.33 | 0.87 |
| Reducing sugar | 12 | 399 | FD+SNV | 4104~7251.4,8825.1~9612,10398.8~11185.7 | 2.65 | 1.34 | 0.42 | 0.91 | | 0.51 | 0.86 | 2.64 | | 106 | 0.61 | 0.89 |
| Sucrose | 18 | 396 | FD+SNV | 4104~ 7251.4,8038.3~ 10398.8 | 49.25 | 5.13 | 1.59 | 0.91 | | 2.39 | 0.78 | 2.14 | | 101 | 1.87 | 0.89 |
| Total sugar | 6 | 367 | FD | 4890.8~ 5677.7,6464.5~ 7251.4,8038.3~ 10398.8 | 51.82 | 5.20 | 1.99 | 0.86 | | 2.11 | 0.83 | 2.46 | | 102 | 1.91 | 0.88 |
| Residues | 10 | 374 | SNV | 4242.8~5276.6,5793.4~7344,7860.8~8377.7 | 48.04 | 5.18 | 1.85 | 0.88 | | 2.02 | 0.85 | 2.57 | | 100 | 1.81 | 0.89 |
| Ratio | | | | | | | | | | | | | | | | |
| Sug/Res | 13 | 399 | FD | 3996~11988 | 1.09 | 0.21 | 0.08 | 0.86 | | 0.10 | 0.75 | 2.02 | | 104 | 0.09 | 0.82 |
| Suc/Total | 14 | 433 | FD | 4242.8~7860.8,8886.8~9403.7 | 0.95 | 0.03 | 0.01 | 0.87 | | 0.01 | 0.81 | 2.27 | | 110 | 0.01 | 0.89 |
| Fru/Glc | 13 | 388 | SNV | 4759.7~5276.6,5793.4~9411.4 | 2.00 | 0.57 | 0.22 | 0.85 | | 0.30 | 0.72 | 1.90 | | 96 | 0.23 | 0.81 |
| Dry weight(Online) | | | | | | | | | | | | | | | | |
| Sugars | | | | | | | | | | | | | | | | |
| Glucose | 16 | 290 | FD | 4890.8~11185.7 | 0.89 | 0.49 | 0.19 | 0.86 | | 0.25 | 0.74 | 1.98 | | 117 | 0.21 | 0.81 |
| Fructose | 20 | 272 | FD | 4104~8038.3,8825.1~11972.5 | 1.60 | 0.53 | 0.23 | 0.83 | | 0.30 | 0.68 | 1.76 | | 104 | 0.27 | 0.79 |
| Reducing sugar | 31 | 285 | FD | 4242.8~9411.4 | 2.44 | 0.97 | 0.33 | 0.90 | | 0.52 | 0.71 | 1.86 | | 116 | 0.46 | 0.84 |
| Sucrose | 10 | 382 | FD+SNV | 5677.7~10398.8,11185.7~11972.5 | 49.59 | 4.83 | 1.86 | 0.86 | | 2.03 | 0.82 | 2.38 | | 111 | 2.35 | 0.82 |
| Total sugar | 8 | 396 | FD+SNV | 5677.7~7251.4,8038.3~11972.5 | 52.31 | 4.84 | 1.95 | 0.84 | | 2.09 | 0.81 | 2.31 | | 112 | 2.36 | 0.80 |
| Residues | 13 | 390 | FD | 4752~6834.8,7853.1~9411.4 | 47.40 | 4.88 | 1.75 | 0.88 | | 1.90 | 0.85 | 2.56 | | 108 | 1.99 | 0.86 |
| Ratio | | | | | | | | | | | | | | | | |
| Sug/Res | 22 | 332 | FD | 4104~11972.5 | 1.11 | 0.20 | 0.06 | 0.91 | | 0.08 | 0.86 | 2.63 | | 131 | 0.07 | 0.88 |
| Suc/Total | 19 | 280 | FD | 4752~9411.4 | 0.95 | 0.02 | 0.01 | 0.88 | | 0.01 | 0.77 | 2.07 | | 110 | 0.01 | 0.81 |
| Fru/Glc | 27 | 282 | FD | 4242.8~7344,7853.1~9411.4 | 1.92 | 0.53 | 0.18 | 0.90 | | 0.26 | 0.77 | 2.09 | | 114 | 0.25 | 0.83 |
| Fresh weight(Online) | | | | | | | | | | | | | | | | |
| Moisture | 22 | 456 | FD+SNV | 4104~ 11972.5 | 73.24 | 2.36 | 0.45 | 0.97 | | 0.55 | 0.95 | 4.30 | | 114 | 0.63 | 0.96 |
| Sugars | | | | | | | | | | | | | | | | |
| Glucose | 31 | 295 | FD | 4104~11972.5 | 0.25 | 0.16 | 0.04 | 0.93 | | 0.07 | 0.78 | 2.13 | | 117 | 0.05 | 0.87 |
| Fructose | 18 | 289 | FD+SNV | 4104~11972.5 | 0.45 | 0.18 | 0.07 | 0.87 | | 0.08 | 0.81 | 2.31 | | 116 | 0.08 | 0.82 |
| Reducing sugar | 24 | 287 | FD | 4242.8~9411.4 | 0.67 | 0.28 | 0.10 | 0.89 | | 0.13 | 0.79 | 2.19 | | 116 | 0.12 | 0.81 |
| Sucrose | 22 | 413 | FD+MSC | 4104~11972.5 | 13.32 | 1.93 | 0.54 | 0.93 | | 0.67 | 0.88 | 2.89 | | 110 | 0.63 | 0.92 |
| Total sugar | 16 | 446 | FD | 5276.6~ 7344,7853.1~ 8377.7 | 14.05 | 2.00 | 0.61 | 0.91 | | 0.68 | 0.89 | 2.96 | | 112 | 0.62 | 0.93 |
| Residues | 11 | 420 | FD+SNV | 5276.6~9411.4 | 12.60 | 1.47 | 0.58 | 0.85 | | 0.62 | 0.82 | 2.38 | | 112 | 0.63 | 0.88 |
N, sample number; SCM, scatter correction methods; SD, standard deviation of reference value; RMSEC, root mean square error of calibration; R^2^, determination coefficient; RMSECV, root mean square error of cross validation; R^2^cv, determination coefficient of cross validation; RMSEP, root mean square error of external validation; R^2^ev, determination coefficient of external validation; RPD, ratio performance deviation; SNV, standard normal variate; MSC, standard multiple scatter; FD, first derivative; FD+SNV, a combination of FD and SNV; FD+MSC, a combination of FD and MSC.
